# RESTORING GLUTAMATE RECEPTOR SIGNALING IN PANCREATIC ALPHA CELLS RESCUES GLUCAGON RESPONSES IN TYPE 1 DIABETES

**DOI:** 10.1101/2021.08.11.455935

**Authors:** Julia K. Panzer, Alejandro Tamayo, Alejandro Caicedo

**Author notes:** Correspondence to: Julia K. Panzer or Alejandro Caicedo, Address: Department of Medicine, University of Miami Miller School of Medicine, 1580 NW 10^th^ Ave, Miami Fl 33136, USA.

## Abstract

Glucagon secretion from pancreatic alpha cells is crucial to prevent hypoglycemia. For reasons still unknown, people with type 1 diabetes lose this glucoregulatory mechanism and are susceptible to dangerous hypoglycemia. Here we show that alpha cells in living pancreas slices from donors with type 1 diabetes failed to secrete glucagon in response to decreases in glucose concentration, thus mirroring the *in vivo* unresponsiveness to hypoglycemia. Glucagon content and responses to KCl depolarization were not affected, suggesting that alpha cells retained their secretory potential. By contrast, alpha cells had severely impaired signaling via glutamate receptors of the AMPA/kainate type. Under healthy conditions, activating these receptors was required to elicit full glucagon responses to decreases in glucose levels. In type 1 diabetes, reactivating residual glutamate receptor function with the positive allosteric modulators cyclothiazide and aniracetam restored glucagon secretion in response to hypoglycemia. These positive allosteric modulators are already approved to treat other conditions and could be repurposed to prevent hypoglycemia and improve management of diabetes.

## MAIN

The alpha cell of the pancreatic islet secretes glucagon to promote gluconeogenesis and glycogenolysis in target organs such as the liver, thus rising blood glucose levels. Although it is now known to be involved in several homeostatic circuits, a very important and clinically relevant function of the alpha cell is to secrete glucagon in response to a drop in glycemia. Glucagon plays a preeminent role in a concerted physiological response named glucose counterregulation that efficiently prevents life-threatening hypoglycemia ^1^. As a result, hypoglycemia is rare in healthy people. For reasons still unknown, in type 1 diabetes, and to some extent in type 2 diabetes, the alpha cell is not capable of regulating its secretion in response to changes in glycemia ^2^. This puts people with diabetes at risk for dangerous hypoglycemia if they are treated with excessive insulin. Understanding the pathophysiological mechanisms that make alpha cells unresponsive to hypoglycemia is thus important to improve the management of diabetes.

As insulin secreting beta cells progressively succumb to the autoimmune attack in type 1 diabetes, alpha cells lose inhibitory input from beta cells. Not surprisingly, basal plasma glucagon levels increase in diabetes, thus exacerbating hyperglycemia ^3^. In spite of this increased basal secretory activity, alpha cells cannot respond adequately to decreases in glycemia ^2^. Recent studies showed that Ca^2+^ channel activity and gene expression decrease in alpha cells in type 1 diabetes, suggesting that the cell may partially lose its membrane excitability ^4,5^. In addition to the electrophysiological properties that allow alpha cells to respond to a drop in glycemia ^6^, a full glucagon response to hypoglycemia relies on autocrine positive feedback mediated by extracellular glutamate acting on ionotropic glutamate receptors of the AMPA/kainate type ^7-11^. These receptors display strong desensitization that is reversed only if glutamate levels subside ^12^. Because glutamate is co-secreted with glucagon ^8^, it is likely that the high basal secretory activity typical of alpha cells in diabetes prolongs the exposure to high levels of glutamate and diminishes glutamate receptor signaling.

Here we tested the hypothesis that the autocrine positive feedback glutamate provides in alpha cells is defective in type 1 diabetes, thus impairing the glucagon counterregulatory response. We used living pancreas slices from non-diabetic donors and donors with type 1 diabetes to determine alpha cell responses to (a) changes in glycemia, (b) agonists, antagonists, and positive allosteric modulators of glutamate receptors, and (c) reference stimuli such as adrenaline and KCl depolarization. We further performed *in vivo* studies in a mouse model of beta cell ablation and *in vitro* studies in living pancreas slices from donors with type 1 diabetes to demonstrate that defective glucagon secretion can be restored by reactivating the glutamate feedback loop in alpha cells.

## RESULTS

It is difficult to study islets in people with type 1 diabetes because there is a limited availability of tissues from donors and conventional techniques may not be adequate to study the obtained islet samples. To overcome these limitations, we used living pancreas slices, which allow functional assessments of damaged and infiltrated islets within their native environment ^13,14^. We examined pancreas slices from 6 nondiabetic donors and 6 donors with type 1 diabetes, which we received from the Organ Processing and Pathology Core (OPPC) of the nPOD program at the University of Florida, Gainesville (Table S1).

### Alpha cells lose their glucose responsiveness in type 1 diabetes

We first investigated if living slices recapitulated known *in vivo* features of islet destruction and dysfunction in type 1 diabetes. Islets in slices from donors with type 1 diabetes displayed a severe loss of beta cells compared to nondiabetic donors (Figure 1a). In line with these results, secretory responses were substantially impaired (Figures 1 b-1f). In stark contrast to slices from nondiabetic donors, slices from donors with type 1 diabetes did not show increases in insulin secretion when stimulated with glucose (17 mM) and only minor responses to KCl depolarization (30 mM; Figures 1b and 1c). Although islets from donors with type 1 diabetes were composed mainly of alpha cells, their functional responses were severely altered. Baseline glucagon secretion was significantly increased (Figures 1d and 1e). When normalized to baseline secretion, the secretory profile revealed a complete loss of glucagon secretion in response to a lowering in glucose concentration (17 mM to 1mM, Figure 1f). However, the secretory capacity of alpha cells was not affected, as seen by the responses to KCl depolarization (Figures 1d and 1f). Glucagon content of lysed tissue slices was not different between the groups (Figure 1g).

**Figure 1:**
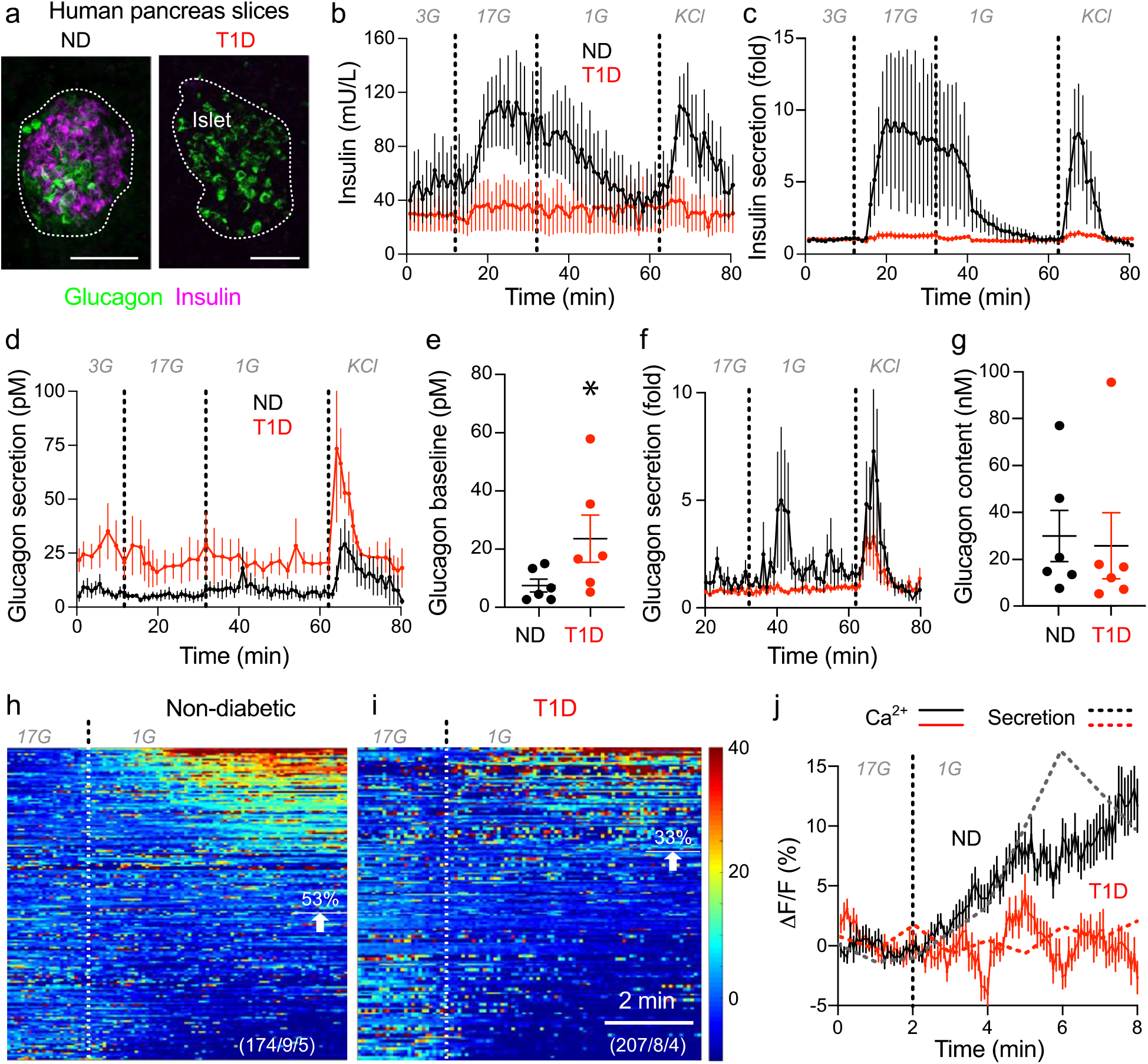
Alpha cells lose their glucose responsiveness in type 1 diabetes. (a) Z stack of confocal images of human tissue slices from a nondiabetic donor (ND) and an individual with type 1 diabetes (T1D) showing beta cells (insulin, magenta) and alpha cells (glucagon, green). Dotted line represents the islet. Scale bar 50 μm. (b) Dynamic insulin secretion of human pancreatic tissue slices from nondiabetic donors (ND, n=6) and individuals with type 1 diabetes (T1D, n=6) during perifusion expressed as absolute insulin (mU/L). (c) Data shown in (b) expressed as fold increase to the mean basal secretion at 3G (Insulin fold). (d) Glucagon secretion measured during the same perifusion experiment shown in (b) expressed as absolute glucagon (pM). (e) Average glucagon secretion during the first 10 min of baseline (3G) shown in (d). (f) Data shown in (d) normalized to average baseline secretion (Glucagon fold). (g) Glucagon content of all perifused slices after secretion experiment expressed as glucagon (nM). (h-i) Heat maps showing in vitro Ca^2+^ dynamics of alpha cells expressed as the fluorescent intensity normalized to the basal signal intensity at 17 mM glucose (17G) at stimulation with low glucose (1G) for (h) 5 nondiabetic donors and (i) 4 individuals with type 1 diabetes. Each row represents a single cell followed over time in the x-axis and their response in magnitude change (%) of the fluorescent intensity over baseline (ΔF/F) shown in the color scale from blue (low intensity) to red (high intensity). For nondiabetic donors cells were sorted by epinephrine response. (j) Average trace of Ca^2+^ dynamics shown in (g and h) compared to average glucagon secretion trace shown in (f) for stimulation with low glucose (1G). Data in (b-d), (f) and (j) are shown as average traces (± SEM). Data in (e) and (g) are shown as dot plots and analyzed with Student’s t test. Asterisks denote significance (*p < 0.05).

To determine the response profiles of individual islet cells we performed imaging of cytoplasmic Ca^2+^ concentration ([Ca^2+^]_i_). We loaded slices with the [Ca^2+^]_i_ indicator Fluo-4 AM and recorded [Ca^2+^]_i_ responses to changes in glucose concentration within the islet by confocal time-lapse imaging (Figures 1h-1i). We only included viable cells that responded to KCl depolarization in our analyses and distinguished alpha cells by their [Ca^2+^]_i_ responses to stimulation with adrenaline ^15^. Changes in fluorescence intensity were calculated as fold-change over baseline intensity and shown as heatmaps that included all recorded cells in slices from nondiabetic donors (Figure 1h) and individuals with type 1 diabetes (Figure 1i). We found that the incidence and magnitude of [Ca^2+^]_i_ responses to a lowering in glucose concentration (17 mM to 1 mM) was diminished in the alpha cell population in donors with type 1 diabetes (Figures 1h-1i). The reduction in the average [Ca^2+^]_i_ response mirrored that of glucagon secretion (Figure 1j). These results indicate that the alpha cell response to a lowering the glucose concentrations is selectively impaired in pancreas slices, which parallels the loss of glucagon counterregulatory responses seen in patients with type 1 diabetes.

### Alpha cell responses to lowering in glucose concentration depend on glutamate receptor activation

To understand what goes awry in diabetes, we first sought to gain a deeper insight into the processes that allow alpha cells to respond to hypoglycemia under non-diabetic conditions. We prepared tissue slices from mice expressing a genetically encoded [Ca^2+^]_i_ indicator (GCaMP3) in alpha cells to study the responsiveness of the whole alpha cell population to changes in glucose concentration and other stimuli (Figure 2; Figures S1 and S2). Surprisingly, only a subset of alpha cells (∼37%) responded to a lowering in glucose concentration to hypoglycemic conditions (7 mM to 1 mM, Figure 2a). A similar proportion of alpha cells (∼31%) showed [Ca^2+^]_i_ responses when we lowered glucose levels from high glucose to normoglycemia (17 mM to 7 mM; Figure 2b). The magnitude and incidence of [Ca^2+^]_i_ responses to drops in glucose concentration were smaller than those elicited by adrenaline (10 μM), kainate (100 μM) and glutamate (100 μM; Figures 2c-2e), all known to be selective and strong alpha cell stimuli ^8,16^. Most of the glucose responsive population was also the one that responded the strongest to kainate, an agonist of ionotropic glutamate receptors (iGluRs) of the AMPA/kainate type.

**Figure 2:**
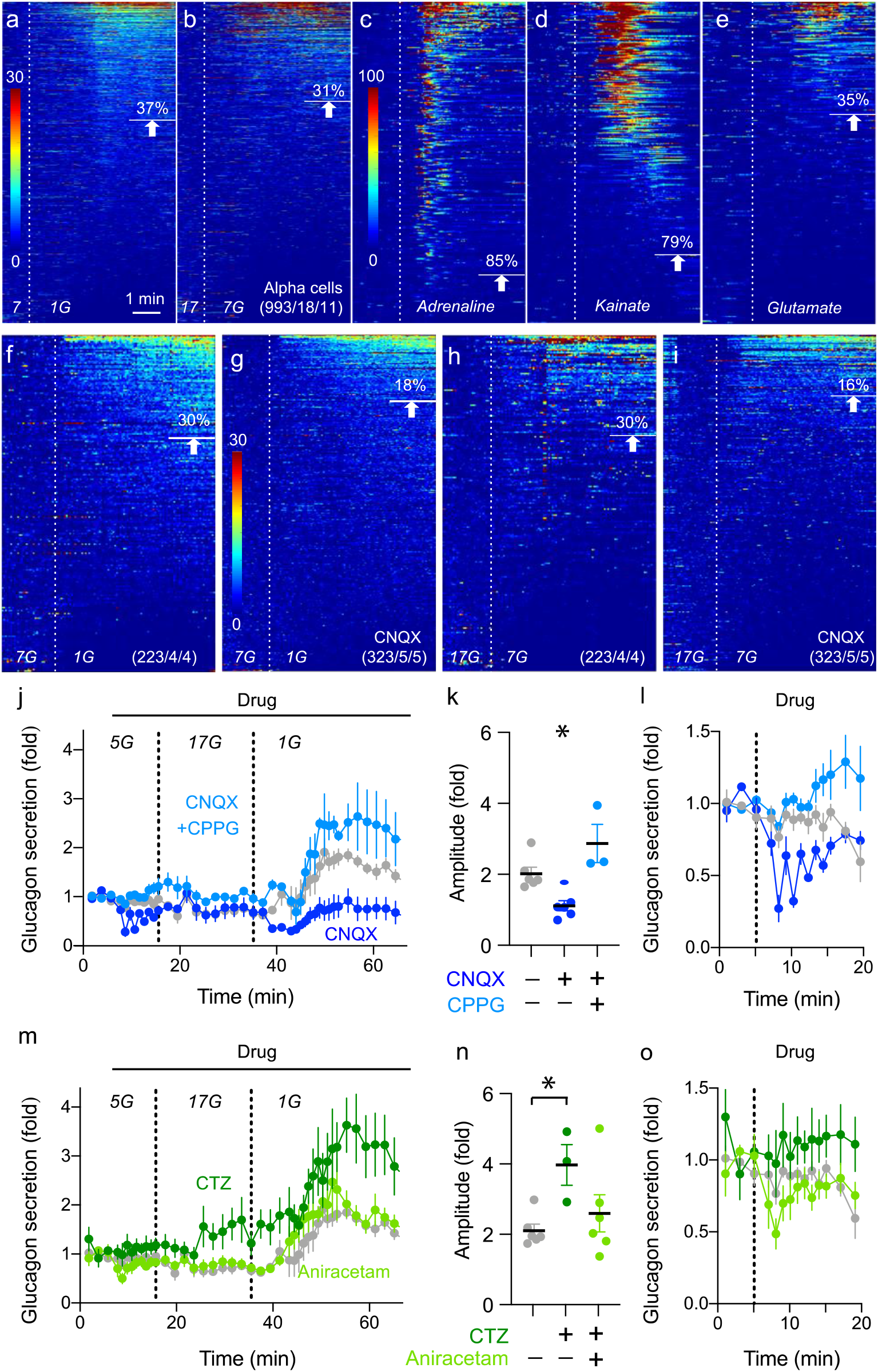
Alpha cell responses to lowering the glucose concentration depend on glutamate receptor activation. (a-i) Heat maps showing in vitro Ca^2+^ responses of alpha cells from Gcg-Cre-GCaMP3 mice within tissue slices. Responses for lowering in glucose concentration (a) from 7 mM to 1 mM and (b) from 17 mM to 7 mM. Alpha cell responses to stimulation with (c) 10 μM adrenalin, (d) 100 μM kainate and (e) 100 μM glutamate. Responses shown for lowering in glucose concentration (f and g) from 7 mM to 1 mM and (h and i) from 17 mM to 7 mM (f and h) in the absence or (g and i) presence of 100 μM CNQX. Responses are expressed as the fluorescent intensity over baseline of GCaMP3 signal. Each row shows a single cell followed over time in the color scale from low intensity (blue) to high intensity (red) expressed as change in magnitude (%). Numbers of cell/islets/animals are indicated in each heatmap. (j) Glucagon secretion from isolated mouse islets during dynamic perifusion in response to changing glucose concentrations and in the presence of 100 μM CNQX or a combination of 100 μM CNQX 100 μM and CPPG. (k) Quantification of glucagon amplitudes shown in (j) during 1 mM stimulation. (l) Magnification of the first 20 min shown in (j). (m) Glucagon secretion from isolated mouse islets during dynamic perifusion in response to changing glucose concentrations and in the presence of 100 μM CTZ or 100 nM Aniracetam. (n) Quantification of glucagon amplitudes shown in (m) during 1 mM stimulation. (o) Magnification of the first 20 min shown in (m). Data in (j), (l), (m) and (o) are shown as average trace (± SEM) normalized to baseline secretion at 7 mM glucose of 150 islets each (control n=6, CNQX n=6, CNQX+CPPG n=3, CTZ n=3, Aniracetam n=6). Data in (k) and (n) are shown as dot plot (± SEM) and analyzed by one-way ANOVA multiple comparisons. Asterisks denote significance (*p < 0.05)

Alpha cells secrete glutamate together with glucagon and express iGluRs of the AMPA/kainate type, establishing an autocrine feedback loop potentiating glucagon secretion ^8^. To determine how alpha cells respond to decreases in glucose concentration depends on this feedback loop, we challenged these responses with the AMPA/kainate receptor antagonist CNQX (100 μM; Figure 2f-2l). The incidence in the alpha cell population of [Ca^2+^]_i_ responses to lowering glucose concentration from 7 mM to 1 mM or 17 mM to 7 mM diminished in the presence of CNQX (Figures 2f-2i). The effects on glucagon secretion were even stronger: glucagon secretion in response to a drop in glucose from 17 mM to 1 mM disappeared in the presence of CNQX (Figures 2j-2k). CNQX also diminished glucagon secretion at basal glucose concentrations (5 mM), suggesting that AMPA/kainate receptors were already activated under normoglycemic conditions (Figure 2l).

The metabotropic glutamate receptor 4 (mGluR4) has also been implicated in glucagon secretion ^17^. Strikingly, the inhibition CNQX produced on glucagon secretion was reversed in the presence of CPPG (100 μM), a potent and competitive antagonist of group II and III metabotropic glutamate receptors (Figures 2m-2o) ^18^. These results suggest that glutamate receptor signaling has major effects on glucagon secretion and involves at least two different receptor types whose activation trigger different signaling pathways (membrane depolarization *versus* decreasing intracellular cAMP levels) that produce opposite effects on secretion.

In view of their therapeutic potential, we tested two positive allosteric modulators of AMPA/kainate receptors, namely cyclothiazide, a compound approved by the FDA to treat edema (Application numbers N018173 and N013157), and aniracetam, a drug available over-the-counter in the US used for its cognition-enhancing effects ^19^. These compounds essentially eliminate rapid desensitization of the AMPA receptor ^20,21^. Cyclothiazide (100 μM) potentiated glucagon secretion from isolated mouse islets when stimulated with low glucose, while aniracetam did not (100 nM; Figures 2m and 2n). Both cyclothiazide and aniracetam amplify AMPA receptor responses by inhibiting receptor desensitization, but aniracetam also potentiates metabotropic glutamate receptor activity ^22^. This effect could be observed as a drop in glucagon secretion (Figure 2o) and might have counteracted the positive effects on glucagon secretion. Nevertheless, our results suggest that potentiating the autocrine glutamate feedback loop can increase glucagon release during conditions of low glucose concentration.

### Alpha cells show defective glucagon counterregulatory responses 2 weeks after beta cell ablation

A main feature of type 1 diabetes is that the autoimmune attack works with surgical precision to exclusively destroy beta cells ^23^. The defect in glucagon secretion seems to track with the progressive loss of beta cell function ^24^. To investigate how alpha cell function changes when beta cells are destroyed, we injected mice with the beta cell toxin streptozotocin. At two weeks after streptozotocin injection, this mouse model mimicked the defective glucagon response typical of type 1 diabetes, both *in vivo* and *in vitro* (Figures 3a-3e). The counterregulatory response to hypoglycemia was defective, most likely because mice did not mount a glucagon response to lowering glucose levels (Figures 3a, 3b, 3d, and 3e; Figure S3). Fasting plasma glucagon concentrations were significantly increased (Figure 3c). Thus, with respect to glucagon secretion, this model reproduced most *in vivo* features of type 1 diabetes ^2,3^ and our *in vitro* results in human slices (Figure 1).

**Figure 3:**
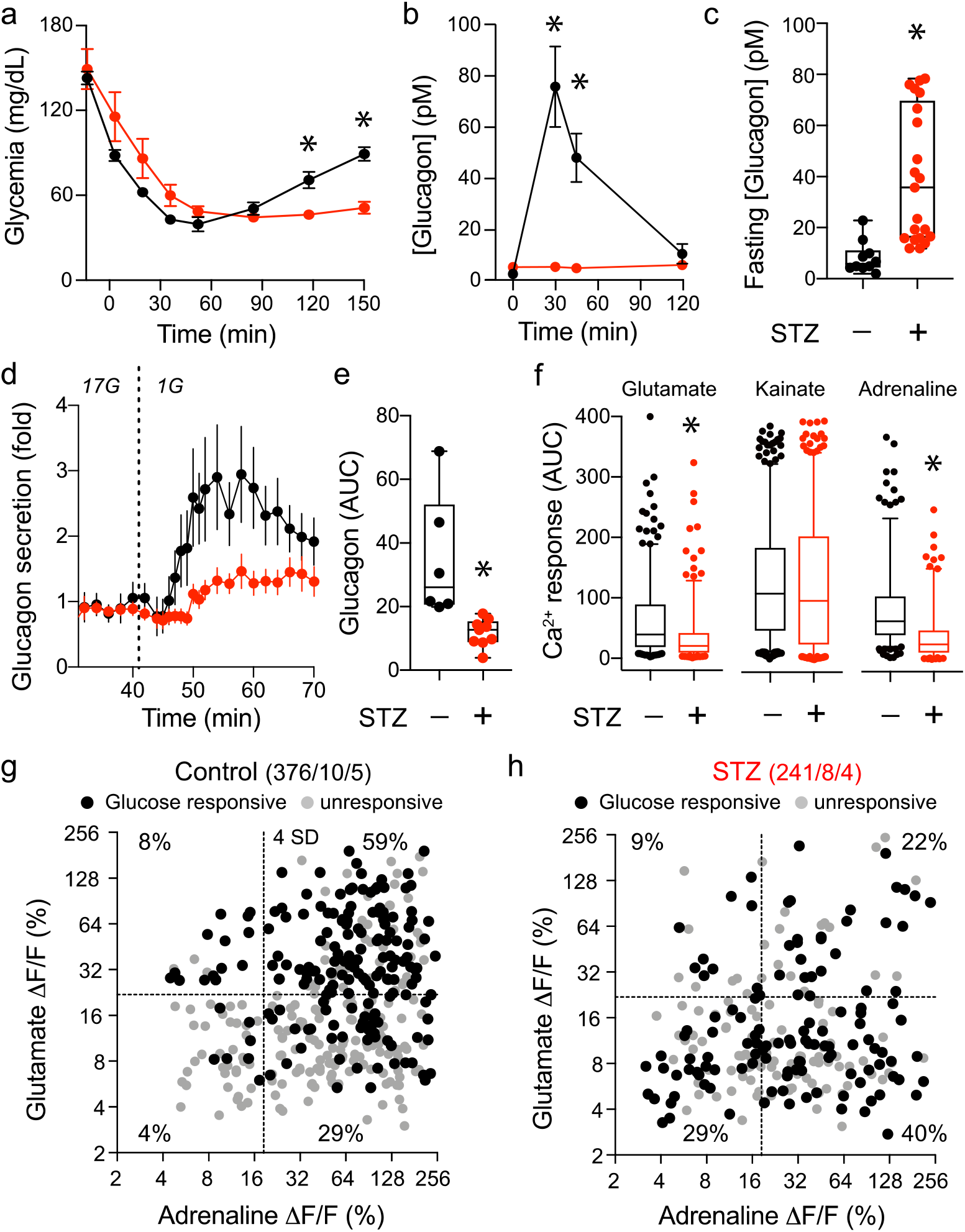
Alpha cells show defective glucagon counterregulatory responses 2 weeks after beta cell ablation. (a and b) Intraperitoneal insulin tolerance test (0.75 U/kg) performed in fasted control (n=30) and rendered diabetic C57Bl6J mice (n=21) 2 weeks after intravenous streptozotocin (STZ, 200mg/kg) injection. (a) Blood glucose excursions and (b) plasma glucagon levels after injection. (c) Quantification of fasted blood glucagon prior to ITT shown in (b) for control (n=10) and STZ (n=21) animals. (d-e) In vitro glucagon secretion from isolated islets of control (n=6) and STZ treated animals (n=9). (d) Average trace of secreted glucagon after lowering glucose from 17 mM to 1 mM normalized to baseline activity and (e) quantification as area under the curve (AUC). (f) Quantification of Ca^2+^ responses from individual alpha cells within slices from Gcg-Cre;GCaMP3 control (n= 376) and STZ treated (n= 241) mice. (g-h) Scattered blots showing maximum amplitudes (%) as fluorescent intensity over baseline (ΔF/F) of individual glucose responsive cells (black dots) and glucose unresponsive cells (grey dots) to stimulation with glutamate and adrenalin for (g) control animals (376 cells, 10 islets, n=5) and STZ treated mice (241 cells, 8 islets, n=4). Percentages represent fraction of glucose responsive alpha cells for each stimulus response. Data in (a), (b) and (c) are shown as average trace (± SEM). Data in (c) and (e) is shown as box-and-whisker plots and compared with Student’s t tests. Data in (f) is shown as box-and-whisker plots of 90 percentile and analyzed with Student’s t test. Asterisks denote significance (*p < 0.05)

We then used [Ca^2+^]_i_ imaging to study how the response pattern of the population of individual alpha cells changed in the streptozotocin model. [Ca^2+^]_i_ responses to a lowering in glucose concentration were reduced in some mice (Figures S3), but the effect was not consistent as it was in human slices (Figure 1) and not as striking as the reduction in glucagon secretion (Figures 3d and 3e). However, we found substantial decreases in the magnitude and number of responses to glutamate and adrenaline (Figures 3f-3h). Responses to glutamate were particularly diminished, suggesting that glutamate receptor signaling was disrupted, which could potentially impact the autocrine alpha cell loop mediated by glutamate.

### Targeting glutamate receptors *in vivo* improves glucagon secretion in response to hypoglycemia in the streptozotocin mouse model

We next determined the effects of the positive allosteric modulators cyclothiazide and aniracetam on glycemia and glucagon secretion *in vivo* using the mouse model. Cyclothiazide (30 mg/kg; i.p.) and aniracetam (30 mg/kg; i.p.) did not alter plasma glucagon levels under fed conditions in healthy mice or in streptozotocin-treated mice fasted to reach normoglycemic conditions (Figures 4a-4d). By contrast, the potent AMPA/kainate agonist kainate increased glucagon secretion in both groups (10 mg/kg, i.p.; Figures 4b and 4d). Of note, when kainate activated glucagon secretion in healthy mice under normoglycemic conditions it decreased glycemia (Figure 4a), but it accelerated the return to hyperglycemia in the diabetic mice (Figure 4c). Thus, activating alpha cells had an impact on glycemia depending on whether or not the islet contained beta cells. The results further indicate that under normoglycemic conditions the positive allosteric modulators did not stimulate glucagon secretion or affect glycemia.

**Figure 4:**
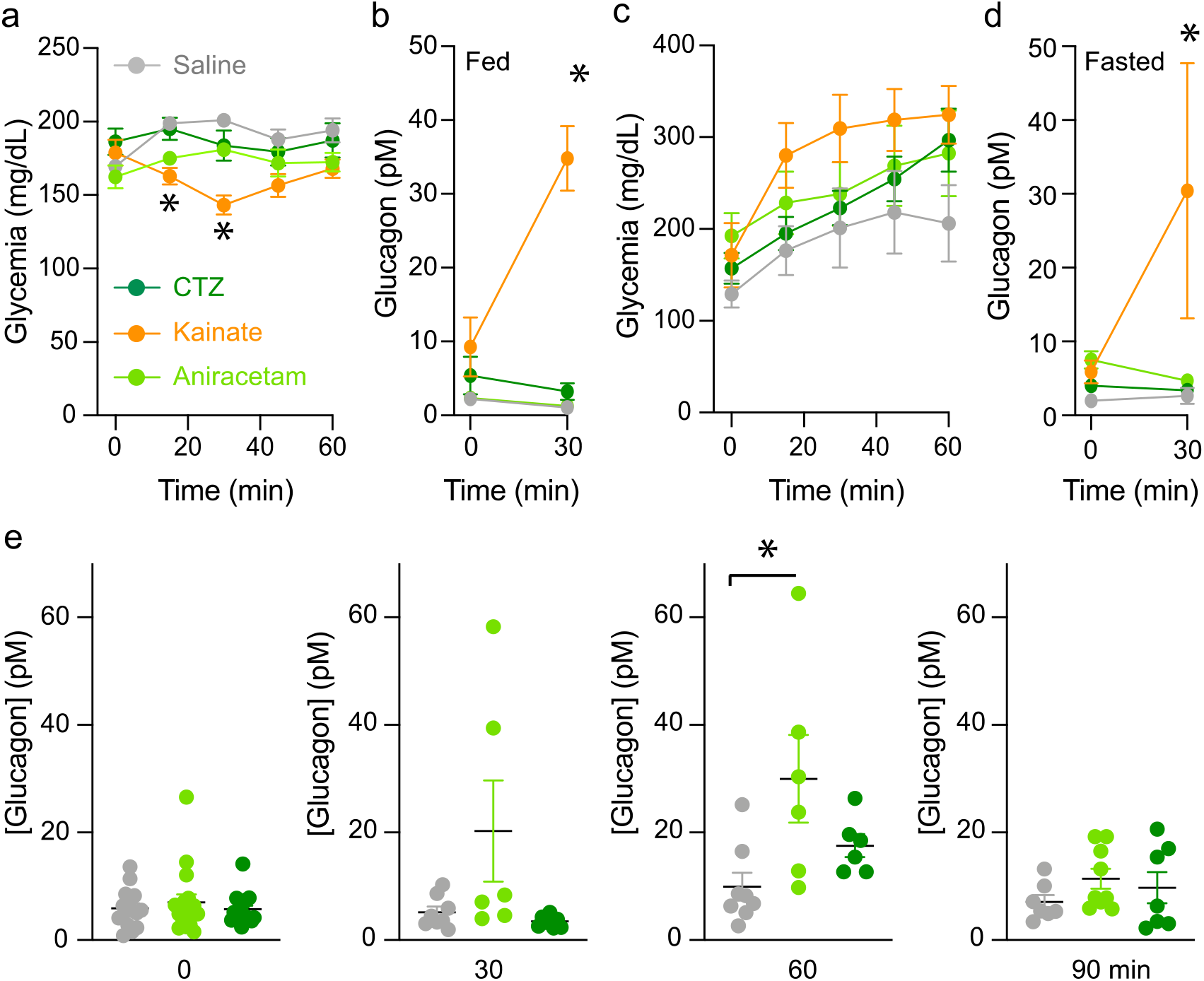
Targeting glutamate receptors *in vivo* improves glucagon secretion in response to hypoglycemia in the streptozotocin mouse model. (a) Non fasting glycemic levels of C57Bl6J mice after injection with saline (n=8), 30 mg/kg CTZ (n=8), 10 mg/kg kainate (n=8) or 30 mg/kg Aniracetam (n =5). (b) Serum glucagon levels of data shown in (a) before and 30 min after injection. (c) Overnight fasted glycemia of rendered diabetic animals with STZ after injection with saline (n=4), 30 mg/kg CTZ (n=9), 10 mg/kg kainate (n=4) or 30 mg/kg Aniracetam (n =9). (d) Serum glucagon levels of data shown in (c) before and 30 min after injection. (e) Plasma glucagon levels of 6 h fasted diabetic animals before injection at 0 min (saline n=15, CTZ n=17, Aniracetam n=17), and 30 min (saline n=8, CTZ n=6, Aniracetam n=6), 60 min (saline n=8, CTZ n=6, Aniracetam n=16) and 90 min (saline n=6, CTZ n=6, Aniracetam n=9) after 0.75 U/kg intraperitoneal insulin injection. Data are shown as average (± SEM) and compared with one-way ANOVA multiple comparisons. Asterisks denote significance (*p < 0.05).

Our *in vitro* results suggest that loss of glucose counterregulation was associated with defective glutamate signaling in alpha cells. To test if reactivating residual glutamate signaling in alpha cells can augment glucagon secretion, we induced hypoglycemia in streptozotocin-treated mice in the presence of the positive allosteric modulators (Figure 4e). Injecting cyclothiazide or aniracetam at the time of the insulin administration increased the glucagon response to hypoglycemia. Although the treatment did not increase glucagon secretion to levels seen in control, healthy mice (Figure 4; ∼ 80 pM *versus* 20-30 pM), these results indicate that glutamate receptors can be potentiated to promote glucagon secretion under hypoglycemic conditions.

### The autocrine feedback loop mediated by glutamate can be potentiated in human alpha cells

To translate the mouse findings to the human islet, we first determined whether the glutamate receptor mechanisms we found in mice were also working in human alpha cells.

Glucagon responses from isolated human islets to a lowering in glucose concentration from 17 mM to 1 mM were potentiated by the positive allosteric modulator cyclothiazide (100 μM; Figures 5a and 5b). These effects were prevented by CNQX (100 μM; Figure 5b). Aniracetam (10 nM) did not amplify the glucagon response, but the combination of aniracetam and CNQX greatly reduced the glucagon response to lowering the glucose concentration (Figures 5d and 5e). This was most likely because CNQX blocked AMPA/kainate receptors, thus unmasking the effects aniracetam also has on the mGluR4 receptor (Figure S4) ^22^, whose activation diminishes glucagon secretion ^17^. Glucagon content was comparable among the groups (Figure 5c and 5f). We could confirm these data in additional living pancreas slices from a non-diabetic donor (Figures 5g and 5h). Here we used a higher concentration of aniracetam (1 mM) because at higher concentrations the effects on AMPA/kainate receptors predominated (Figure S4). As in the mouse, our results indicate that glutamate signaling impacts glucagon secretion from human alpha cells.

**Figure 5:**
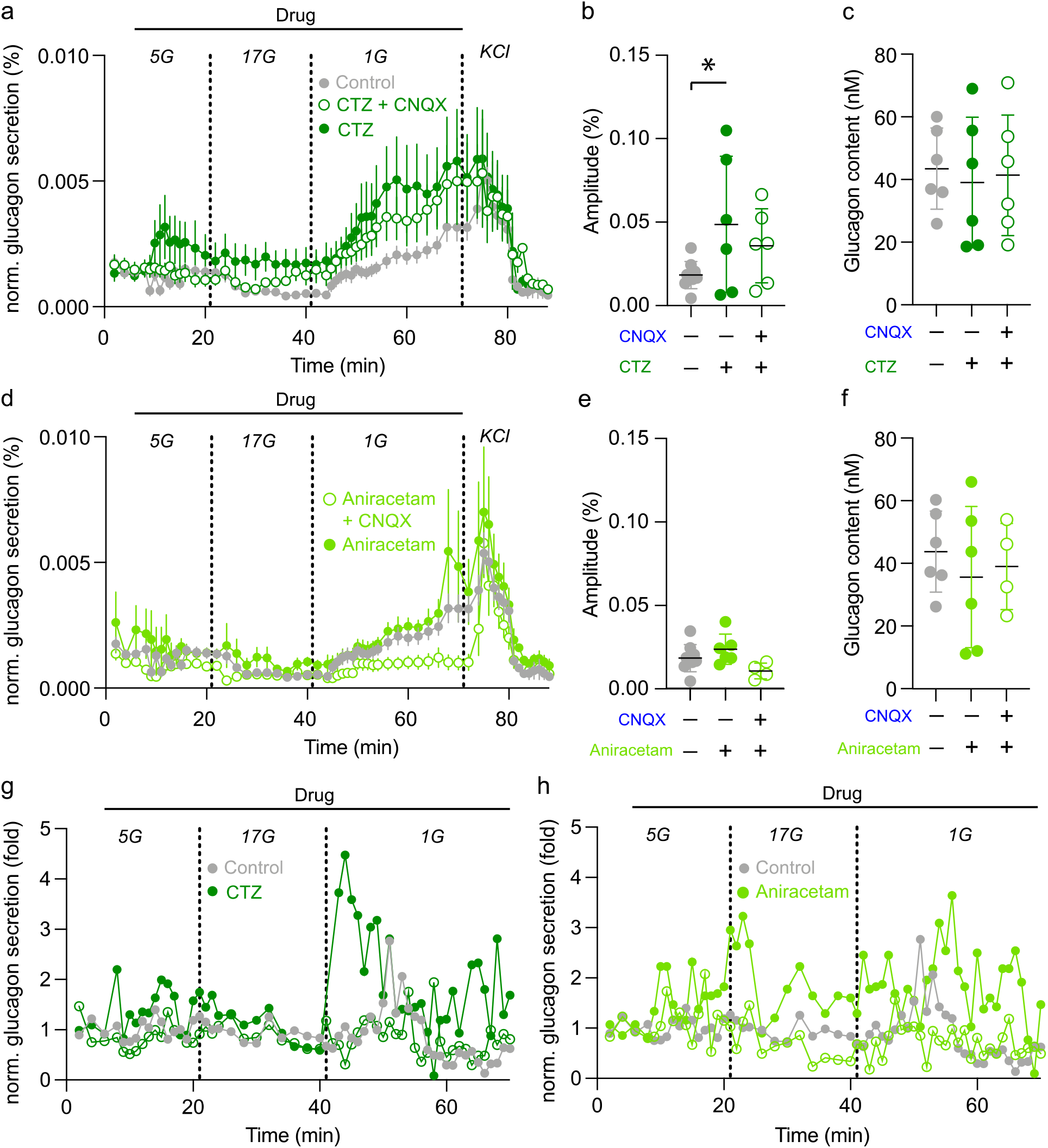
The autocrine feedback loop mediated by glutamate can be potentiated in human alpha cells. (a)Glucagon secretion of isolated human islets in response to changes in glucose concentration. Perifusion experiments performed in the presence of glucose only (n=6), 100 μM CTZ (n=6), and a combination of 100 μM CTZ and 100 μM CNQX (n=6). (b) Quantification of glucagon amplitudes shown in (a) during 1 mM stimulation. (c) Total glucagon content of perifused human islets used in (a). (d) Glucagon secretion of isolated human islets in response to changes in glucose concentration. Perifusion experiments performed in the presence of glucose only (n=6), 10 nM Aniracetam (n=6) or a combination of 10 nM Aniracetam and 100 μM CNQX (n=4). (e) Quantification of glucagon amplitudes shown in (d) during 1 mM stimulation. (f) Total glucagon content of perifused human islets used in (d). (g-h) Glucagon secretion normalized to baseline values of tissue slices from a single nondiabetic human donor in response to changes in glucose concentration in the presence of (g) 100 μM CTZ or a combination of 100 μM CTZ and 100 μM CNQX and (h) in the presence of 1mM Aniracetam or a combination of 1 mM Aniracetam and 100 μM CNQX. Glucose concentrations are indicated above each trace. Average traces (a and d) shown as mean (± SEM) percent of glucagon content from perifused islets. Traces (h and i) are normalized to the mean baseline within the first 6 minutes. Data is analyzed by one-way ANOVA multiple comparisons. Asterisks denote significance (*p < 0.05).

### AMPA/kainate receptor signaling is strongly reduced but can be restored in type 1 diabetes

To investigate how glutamate receptor signaling in alpha cells changes in type 1 diabetes, we perform [Ca^2+^]_i_ imaging in living human pancreas slices loaded with the [Ca^2+^]_i_ indicator Fluo-4 AM. We used responses to adrenaline to identify alpha cells in slices from non-diabetic donors. Responses to adrenaline could not be used to identify alpha cells in donors with type 1 diabetes because these responses were also affected. However, we anticipated (as shown in Figure 1a) that alpha cells in these donors represent the majority of cells exhibiting increases in [Ca^2+^]_i_. All cells were further preselected by their response to KCl depolarization (4 × SD of baseline activity) to ensure cell viability. Ca^2+^ responses to glutamate and kainate could be readily elicited in islet cells in slices from non-diabetic donors but were greatly reduced in slices form diabetic donors (Figures 6a-6d). The incidence of responsive cells decreased from 41% to 18% for glutamate and from 69% to 21% for kainate stimulation.

**Figure 6:**
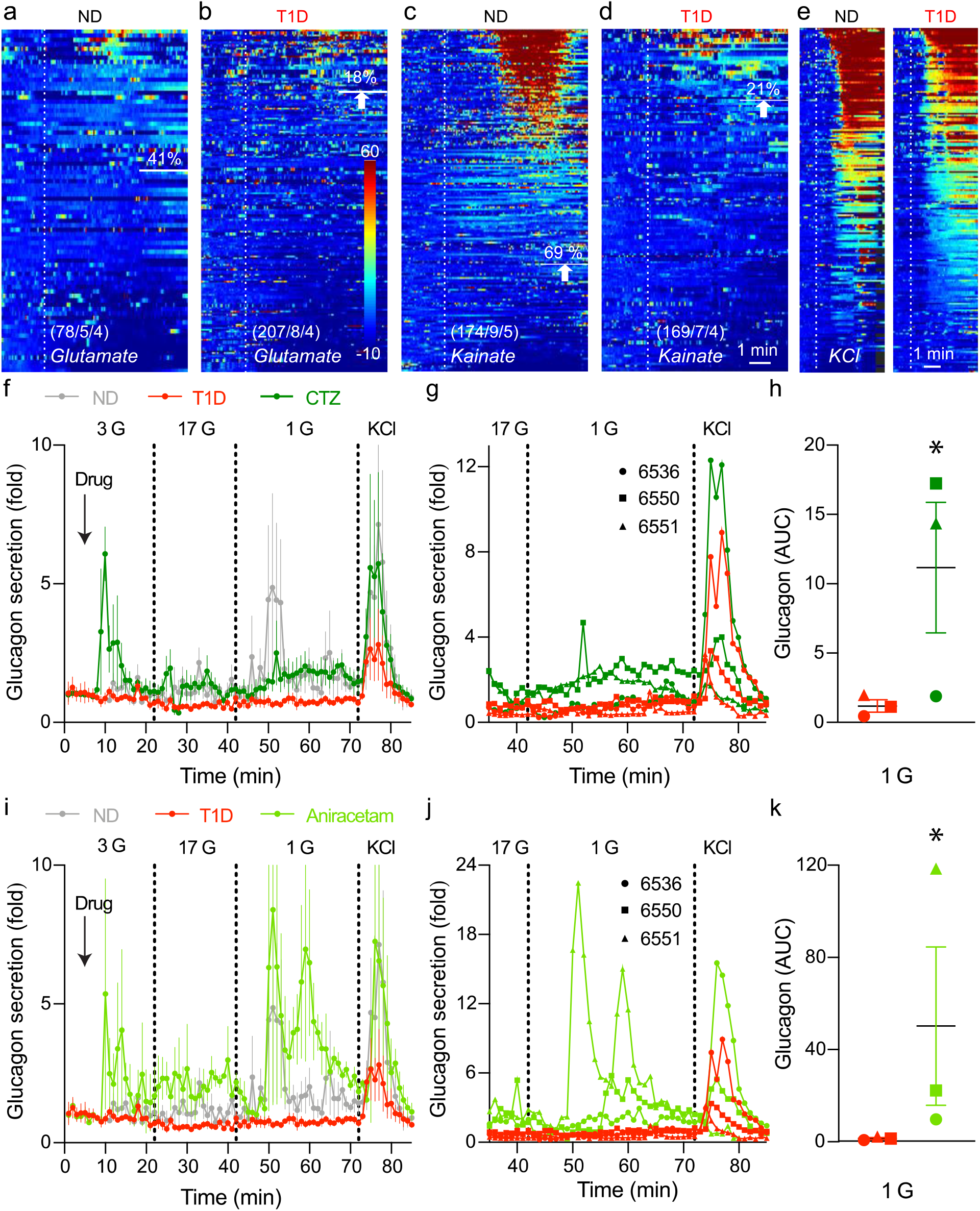
AMPA/kainate receptor signaling is strongly reduced in type 1 diabetes. (a-d) Heat maps showing in vitro Ca^2+^ dynamics of alpha cells expressed as the fluorescent intensity of Fluo4 for (a and c) nondiabetic donors and (b and d) individuals with type 1 diabetes. Each row represents a single cell followed over time in the x-axis and their response in magnitude change (%) of the fluorescent intensity over baseline (ΔF/F) activity. Alpha cell responses (a and b) to 100 μM glutamate and (c and d) 100 μM kainate. Numbers of cell/islets/donors are indicated in each heat map. For nondiabetic donors cells were sorted by epinephrine response. (e) Heat maps showing Ca^2^+ responses to membrane depolarization with 25 mM KCl to ensure cell viability of slices from nondiabetic donors (ND) and individuals with type 1 diabetes (T1D). (f) Glucagon secretion of human tissue slices in response to changes in glucose concentration. Perifusion experiments were performed in the presence of 100 μM CTZ (n=3) and are compared to traces from Figure 1(f). (g) Individual traces of glucagon secretion from 3 donors with T1D stimulated with glucose in the presence and absence of 100 μM CTZ. (h) Quantification of glucagon secretion to 1G stimulation shown in (f and g) calculated as area under the curve. (i) Glucagon secretion of human tissue slices in response to changes in glucose concentration. Perifusion experiments were performed in the presence of 1 mM Aniracetam (n=3) and are compared to traces from Figure 1 (f). (j) Individual traces of glucagon secretion from 3 donors with T1D stimulated with glucose in the presence and absence of 1 mM Aniracetam. (k) Quantification of glucagon secretion to 1G stimulation shown in (i and j) calculated as area under the curve. Traces are normalized to the mean baseline within the first 6 minutes. Traces in (f) and (i) are shown as mean (± SEM) with n=6 for non-diabetic donors, n=6 for donors with T1D and n=3 for donors with T1D stimulated with either CTZ or Aniracetam.

There was a residual response to glutamate receptor agonists in a subset of alpha cells in type 1 diabetes (responders in Figure S5). We therefore tested if this response could be potentiated to rescue the alpha cell response to hypoglycemia. We exposed slices from three donors with type 1 diabetes to cyclothiazide (100 μM, Figure 6f-6h) or aniracetam (1 mM; Figure 6i-6k) and stimulated alpha cells with a drop in glucose from 17 to 1 mM. Our results show that both potentiators increased the glucagon response to low glucose in all cases (Figures 6h and 6k). These drugs did affect residual insulin secretion (Figure S6).

## DISCUSSION

Our study establishes that the alpha cell is not able to mount an efficient glucagon response to hypoglycemic conditions if glutamate receptor signaling is deficient. Inhibiting glutamate receptors of the AMPA/kainate type strongly reduced Ca^2+^ and glucagon responses to decreases in glucose concentration. In type 1 diabetes, glutamate receptor signaling in alpha cells is severely impaired, thus preventing the positive feedback glutamate provides for glucagon secretion. The main conclusion from our studies is that alpha cells in people with type 1 diabetes lose this important autocrine mechanism that guarantees a glucagon counterregulatory response. Our results further demonstrate that residual glutamate signaling can be potentiated with positive allosteric modulators to rescue glucagon secretion.

Thanks to the nPOD initiative to procure and characterize pancreases from cadaveric organ donors with type 1 diabetes, we were able to study the physiology of pancreatic islets in rare specimens from type 1 diabetic donors. This allowed us to determine that the deficiencies commonly reported for glucagon secretion in patients with type 1 diabetes can be translated to pancreatic slices *in vitro*, as also recently shown for isolated islets ^5^. Strikingly, the main alpha cell impairment appears to be selective. Both *in vivo* and *in vitro*, full glucagon responses can be elicited by stimuli such as arginine, adrenaline, and KCl depolarization, and glucagon content was not reduced, suggesting that alpha cells retain their secretory potential (present study; ^2,5^. By contrast, glucagon responses to decreases in glucose concentration are barely detectable. That this defect is present in isolated islets or pancreatic slices not only implies that the deficiency is intrinsic to the pancreatic islet, but also that pathophysiological mechanisms can be studied *in vitro*, as we did here.

A major result of this study is that glutamate receptors of the AMPA/kainate can barely be activated in alpha cells of people with type 1 diabetes. There may be a general loss of alpha cell excitability ^4^, although responses to KCl depolarization *in vitro* and arginine *in vivo* show otherwise (Brissova *et al*., 2018; Gerich *et al*., 1973). Expression of genes encoding for the glutamate receptors may be decreased ^4,5^. However, given the high basal glucagon secretion we measured in slices and the high plasma glucagon levels prevalent in type 1 diabetes, it is also likely that these receptors do not respond because they are chronically desensitized. In line with the notion that AMPA/kainate receptors are chronically desensitized but still present in alpha cells in type 1 diabetes, we found that glucagon secretion in response to a lowering in glucose concentration could be rescued by the positive allosteric modulators aniracetam and cyclothiazide. These inhibitors of AMPA/kainate receptor desensitization also partially restored glucagon secretion in response to a decrease in glucose concentration in diabetic mice *in vivo*. These effects were context-specific; these potentiators did not affect glycemia in the absence of hypoglycemia (Figure 4a-d).

It is clear that what is missing in type 1 diabetes is the inhibitory influence from beta cells. Indeed, most secretory products from beta cells have been reported to inhibit alpha cells. That alpha cell deficiencies can be recapitulated in the streptozotocin rodent model in the absence of autoimmunity indicates that the absence of beta cells is crucial ^25^. While we cannot rule out intrinsic defects in alpha cell function (e.g., responses to adrenaline were diminished), the abnormal alpha cell response to glucose levels can be attributed in part to a loss of inhibitory signaling. It has been proposed that a cessation of these inhibitory signals from beta cells in the islet during hypoglycemia is a necessary signal for the glucagon response from neighboring alpha cells [the “switch-off” hypothesis; ^25,26^]. AMPA/kainate receptors are known to strongly desensitize in the sustained presence of glutamate, and recovery from desensitization requires the dissociation of glutamate, that is, it needs extracellular glutamate levels to subside ^27^. But this cannot happen because glutamate is likely secreted at high rates together with glucagon. To recover from receptor desensitization, the alpha cell needs to rest and reset. We propose that the inhibition of glucagon and glutamate secretion by the inhibitory signals from beta cells is needed for AMPA/kainate receptors in alpha cells to recover from desensitization, allowing the glutamate autocrine feedback loop to be fully activated by a subsequent drop in glucose levels.

Based on our results, we conclude that in type 1 diabetes alpha cells lose an important mechanism that allows efficient glucagon secretion in the context of glucose counterregulation. We demonstrate that AMPA/kainate receptor signaling has a strong impact on glucagon secretion and that this signaling is impaired in type 1 diabetes. Our findings pave the way for testing the role of the glutamate autocrine feedback loop *in vivo* in clinical studies, in particular because several drugs modulating these receptors are already approved for use in human beings (e.g., cyclothiazide, aniracetam and several other nootropics and ampakines). Studying the effects of these compounds in hypoglycemic clamps could not only reveal the impact of glutamate receptor signaling on glucagon responses, but also provide evidence that these receptors can be targeted to prevent recurrent hypoglycemia in people with type 1 diabetes. We expect our studies to eventually help improve the management of type 1 diabetes.

## ACKNOWLEDGEMENTS

This research was performed with the support of the Network for Pancreatic Organ donors with Diabetes (nPOD; RRID:SCR_014641), a collaborative type 1 diabetes research project supported by JDRF (nPOD: 5-SRA-2018-557-Q-R) and The Leona M. & Harry B. Helmsley Charitable Trust (Grant#2018PG-T1D053, G-2108-04793). The content and views expressed are the responsibility of the authors and do not necessarily reflect the official view of nPOD. Organ Procurement Organizations (OPO) partnering with nPOD to provide research resources are listed at http://www.jdrfnpod.org/for-partners/npod-partners/. The authors are grateful to the donors and their families for their invaluable contribution. We thank Alexander Kao and Guillermo Camarena from Biorep Technologies, Inc. for developing the tissue slice chambers used in this paper. This work was funded by NIH grants R56DK084321 (A.C.), R01DK084321 (A.C.), R01DK111538 (A.C.), R01DK113093 (A.C.), U01DK120456 (A.C.), R33ES025673 (A.C.), R21ES025673 (A.C.), and R01DK130328 (A.C.), the Leona M. and Harry B. Helmsley Charitable Trust grants G-2018PG-T1D034 (A.C.) and G-1912-03552 (A.C.).

## AUTHOR CONTRIBUTIONS

J.K.P. designed and performed physiological experiments with slices/islets and performed immunohistochemistry. A.T. conducted vivo experiments, maintained animal colonies, performed islet isolations and contributed to physiological experiments. J.K.P. and A.C. designed the study, analyzed and interpreted data and wrote the paper. All authors discussed the results and critically edited the manuscript.

## DECLARATION OF INTERESTS

The authors declare no competing interests.

## METHODS

### Mouse models

For in vivo and ex vivo measurements of hormone secretion we used male C57Bl6J (JAX stock #000664) mice, age 10-14 weeks, purchased from The Jackson Laboratory (JAX). After in vivo measurements, the pancreas of these mice was processed for islet isolation as described below. For ex vivo measurements of cytosolic Ca^2+^ in alpha cells, we generated mice that express GCaMP3 selectively in alpha using the Cre-Lox system. Briefly, we crossed female Gcg/tm 1.1(icre)Gkg/J mice (JAX stock #030663), with male mice that express GCaMP3 downstream of a loxP-flanked STOP cassette (JAX stock #029043). Only F1 male mice were used, 8-14 weeks old. The pancreas of these animals was used to generate pancreatic tissue slices as described below. All experiments were conducted according to protocols and guidelines approved by the University of Miami Institutional Animal Care and Use Committee.

The University of Miami complies with the Animal Welfare Act of 1966 (PL89-544) as amended by the Welfare Act of 1970 (PL91-279), adheres to the principals stated in the guide for the care and use of laboratory animals – NIH publication #85-23 (revised) and is accredited by the Association for Assessment and Accreditation of Laboratory Animal Care.

### Human organ donors

Pancreas tissue from human donors with or without T1D was procured by the nPOD program at the University of Florida in Gainesville (https://www.jdrfnpod.org). All procedures were performed according to the established SOPs by the nPOD/OPPC and approved by the University of Florida Institutional Review Board (IRB201600029) and the United Network for Organ Sharing (UNOS) according to federal guidelines, with informed consent obtained from each donor’s legal representative. Demographic data, hospitalization duration, and organ transport time were obtained from hospital records. Donor pancreata were recovered, placed in transport media on ice, and shipped via organ courier to the University of Florida. Donor information is listed in Table S1. Human pancreatic islets were obtained from non-diabetic, cadaveric donors through Prodo Laboratories. Detailed donor information is listed in Table S2.

### Intraperitoneal Insulin-Tolerance Tests

Intraperitoneal insulin-tolerance tests (IPITTs) were performed in mice after 6 h fasting. To induce hypoglycemia, mice were injected with insulin (intraperitoneal 0.75 U/kg) and blood glucose was monitored at predetermined time points after the injection. STZ treated animals were injected with insulin (intraperitoneal 0.75 U/kg) 3 h prior to the ITT to decrease glycemia to comparable levels with control animals and ensure lowering to hypoglycaemic setpoints during the experiment.

### Islet isolation procedure

Pancreatic islets were isolated by enzymatic digestion using collagenase type V from *Clostridium histolyticum* (Sigma), dissolved in Hanks’ Solution (Sigma) to a final concentration of 0.4 mg/mL. Briefly, mice were anesthetized with isofluorane and euthanized by cervical dislocation. 3 mL of cold collagenase solution was injected into the common bile duct. The injected pancreas was transferred in a 50 mL Falcon tube containing 2 mL of collagenase solution stored on ice. Digestion was performed in a water bath at 37°C for 20 min followed by gentle shaking by hand. The reaction was stopped by adding 25 mL ice-cold Hank’s solution. After 1 min centrifugation at 1400 rpm at 8°C, supernatant was discarded, the pellet was resolved in 25 mL Hanks’ solution and passed through a small sterile metal strainer. The solution was centrifuged for 1 min at 1400 rpm at 8°C, resuspended in 25 mL Hanks’ solution and centrifuged as before. The pellet was resuspended in 12.5 mL Histopaque® 1077(Sigma) and 10 mL of Hanks’ solution was added carefully. After 24 min centrifugation at 3510 rpm with minimum acceleration and breaking, supernatant was collected and washed with 45 mL Hanks’ solution. After 1 min centrifugation at 1500 rpm, islet pellet is resuspended in 10 mL CMRL media (Corning). Islets were handpicked and rested overnight in an incubator at 37°C and 5 % CO_2_.

### Preparation of living tissue slicing

Tissue slices were prepared from male Gcg-Cre-GCaMP3 mice (8-14 weeks old). Briefly, mice were anesthetized with isofluorane and euthanized by cervical dislocation. Low gelling temperature agarose (1.2%, Sigma, Cat# A9414, dissolved in HEPES-buffered solution without BSA) was injected in the common bile duct using a 30-gauge needle and 5 mL syringe. After injection, small tissue pieces were cut and embedded further in agarose and placed at 4C for 10 min. Pancreatic slices were then cut (150 mm thickness) on a vibrating blade microtome (VT1000S, Leica) and incubated in HEPES-buffered solution (125 mM NaCl, 5.9 mM KCl, 2.56 mM CaCl2, 1 mM MgCl2, 25 mM HEPES, 0.1% BSA, pH 7.4) containing 5.5 or 7 mM glucose and supplemented with aprotinin (25KIU/mL, Sigma, Cat**#** A1153).

### Dynamic Measurements of hormone secretion

To assess glucose-stimulated hormone release of human pancreatic tissue slices, 3 viable slices were placed into slice perifusion chambers (Biorep Technologies, part No. PERI-CHAMBER) and connected to a perifusion system with automated tray handling (Biorep Technologies, cat. no. PERI4-02-230-FA). For isolated islets (human/mouse), ∼100-150 islets were placed in perifusion columns (Biorep Technologies, part No: PERI-PSC-001) and connected to the perifusion system. Slices/islets were perfused at a flow rate of 100 μL/minute and samples were collected in 96-well plates with a 60 sec interval. Initially tissue samples were flushed for 90 minutes with baseline Hepes buffer to wash out accumulated hormones and enzymes, followed by stimulation with high glucose, low glucose and KCl (with and without the addition of iGluR antagonists/agonists as indicated). Perfusates were stored at −20°C until insulin/glucagon content was measured using commercially available ELISA kits (Mercodia, cat. no. 10-1113-01 and 10-1281-01). Concentrations of insulin/glucagon were obtained by plotting the absorbance of calibrators versus their concentration using sigmoidal four parameter logistic with Prism 9 software (GraphPad software, La Jolla, CA). Data is presented either normalized to average baseline secretion (fold) or as percent of total hormone content (%). Hormone content was measured from lysates obtained from all islets/slices used after the experiment.

### Ca^2+^ imaging

2-3 viable human slices were incubated with Fluo4-AM (6 μM, Invitrogen cat. No. F1221) for 1h in 3 mM HEPES buffer (125 mmol/l NaCl, 5.9 mmol/l KCl, 2.56 mmol/l CaCl_2_, 1 mmol/l MgCl_2_, 25 mmol/l HEPES, 0.1% BSA [wt/vol.], pH 7.4) supplemented with aprotinin (25KIU, Sigma, cat# A1153) at room temperature in the dark. Slices were then placed on a coverslip in an imaging chamber (Warner instruments, Hamden, CT, USA) for imaging on a Leica TCS SP5 upright laser-scanning confocal microscope (Leica Microsystems, Wetzlar, Germany). Slices were continuously perfused with HEPES-buffer solution with glucose and prototypic stimuli. Confocal images were acquired with LAS AF software (Leica Microsystems) using a 40X water immersion objective (NA 0.8). A resonance scanner was used for fast image acquisition producing time-lapse recordings spanning 50-100 μm of the slice (z-step: 5 μm, stack of ten confocal images with a size of 512 × 512 pixels) at 5 sec intervals (XYZT imaging). Backscatter light was collected upon excitation with the 633 nm laser and Fluo4-AM fluorescence was excited at 488 nm and emission detected at 510–550 nm. To quantify changes in intracellular Ca^2+^ levels, we drew regions of interest around individual islet cells and measured changes in mean GCaMP3/Fluo4-AM fluorescence intensity using ImageJ. Regions of interest were drawn using single planes. A detailed description of how Ca^2+^imaging data were analyzed and quantified is provided as supplementary material (Figure S2). Briefly, changes in fluorescence intensity were expressed as percentage changes over baseline (ΔF/F). The baseline was defined as the mean intensity of 3 min baseline prior to each stimulus either as heatmaps using MATLAB R2019a (MathWorks software, Natick, MA) or as average traces using Prism 9 (GraphPad software, La Jolla, CA).

### Immunofluorescent staining

Slices are washed for 10 min in PBS prior to incubation in 1x blocking solution (BioGeneX, cat No. HK085-5K, diluted in PBS 0.3% Triton X-100) for 2 hours at room temperature. Primary antibodies are diluted in 1x blocking solution and incubated with antibodies against insulin (1:2000, guineapig, Dako, cat. no. A-0546), glucagon (1:500, mouse, Sigma, cat. no. G2654) and somatostatin (1:500, rabbit, Millipore, cat. No. MAB354) overnight at 4°C while shaking. Slices were washed three times for minimum 30 minutes with PBS and then incubated with secondary antibodies diluted in PBS overnight at 4°C while shaking. Slices were washed three times for 10 minutes with PBS and kept at 4°C in the dark until imaging.

### Statistical Analyses

For statistical comparisons we used Prism 9 (GraphPad software, La Jolla, CA) and performed Student’s t tests (unpaired) or one-way analysis of variance (ANOVA) corrected for multiple comparisons (using ordinary one-way ANOVA). P-values < 0.05 were considered statistically significant. Throughout the manuscript we present data as mean ± SEM.

## Notes

### Competing Interest Statement

The authors have declared no competing interest.

