## Supplementary figures/tables for "RESTORING GLUTAMATE RECEPTOR SIGNALING IN PANCREATIC ALPHA CELLS RESCUES GLUCAGON RESPONSES IN TYPE 1 DIABETES"

### SUPPLEMENTARY TABLES

**Table S1.**

| nPOD Case ID | Donor type | Age (yrs) | Diabetes duration | AAb status | Gender | Race | BMI | HbA1c | COD |
|---|---|---|---|---|---|---|---|---|---|
| 6502 | ND | 6 | | Negative | Male | Native American | 15.2 | 5.7 | Anoxia |
| 6516 | ND | 21 | | Negative | Male | Caucasian | 28.8 | 5.5 | Head trauma |
| 6531 | ND | 19 | | Negative | Female | Hispanic | 30 | 5.6 | Head trauma |
| 6535 | ND | 31 | | Negative | Female | Caucasian | 29.6 | 5.5 | Head trauma |
| 6537 | ND | 33 | | Negative | Male | Caucasian | 20.8 | 5.6 | Head trauma |
| 6546 | ND | 22 | | Negative | Male | Asian | 23.7 | 5.6 | Anoxia |
| 6523 | T1D | 12 | 3 years | GADA+ mIAA+ | Female | African American | 22.5 | 11.1 | Anoxia |
| 6528 | T1D | 14 | 0 years | Negative | Male | African American | 24 | 12.3 | Anoxia |
| 6533 | T1D | 4 | 0 years | IA2A+ mIAA+ ZnT8A+ | Female | Caucasian | 17.7 | 11.4 | Anoxia |
| 6536 | T1D | 20 | 4 years | GADA+ | Female | Caucasian | 25.4 | 12.7 | Anoxia |
| 6550 | T1D | 25 | 0 years | GADA+ ZnT8A+ | Male | Caucasian | 16 | >14 | DKA |
| 6551 | T1D | 20 | 7 months | | Male | | 23.1 | 6.4 | Anoxia |

**Characteristics of organ donors utilized for tissue slices. Related to Figure 1 and 6.**
Table showing characteristic for nondiabetic donors and organ donors with type 1 diabetes. Organs were received through the nPOD program and slices were generated by the nPOD Organ Processing and Pathology Core [OPPC] and shipped overnight. ND = nondiabetic; T1D = type 1 diabets; AAb = autoantibody; COD = cause of death.

**Table S2.**

| Prodo Case ID | Donor type | Age (yrs) | Gender | Race | BMI | HbA1c | COD |
|---|---|---|---|---|---|---|---|
| HP 21079 | ND | 59 | Male | Caucasian | 22.6 | 5.2 | Stroke |
| HP 21086 | ND | 31 | Female | Caucasian | 25.6 | 5.1 | Stroke |
| HP 21048 | ND | 19 | Male | Caucasian | 23.1 | 5.8 | Head trauma |
| HP 21046 | ND | 39 | Male | Caucasian | 27.6 | 5.5 | Anoxia |
| HP 21115 | ND | 37 | Male | Asian | 28.4 | 5.2 | Head trauma |
| HP 21155 | ND | 18 | Male | Hispanic | 20.4 | 5.4 | Head trauma |
| HP 21189 | ND | 26 | Male | Hispanic | 26.3 | 5.4 | Head trauma |
| HP 21203 | ND | 25 | Male | Hispanic | 26.5 | 5.8 | Head trauma |

**Table S2: Characteristics of organ donors utilized for isolated islets. Related to Figure 1 and 6.**
Table showing characteristic for nondiabetic donors. Islets were obtained through Prodo Laboratories.
ND = nondiabetic. COD = cause of death.

### SUPPLEMENTARY FIGURES

Figure S1

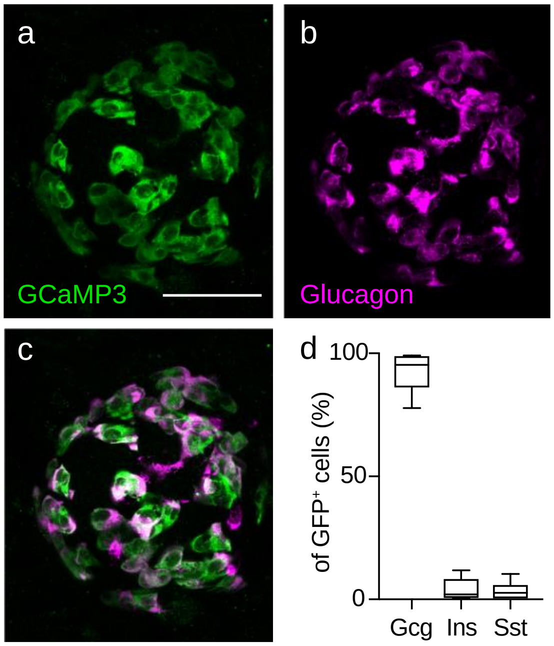


**Figure S1: Recombination efficiency of Gcg-Cre-GCaMP3 mouse model. Related to Figure 2.** (a-c) Z stack of confocal images of an islet within a tissue slice from mice expressing a genetically encoded $Ca^{2+}$ indicator in all alpha cells (Gcg-Cre-GCamP3). (a) GCaMP3 expression (GFP, green), (b) alpha cell staining (glucagon, magenta) and (c) merged image of both. Scale bar 50 µm. (d) Quantification of recombination efficiency shown as percent of GFP positive cells.

Figure S2

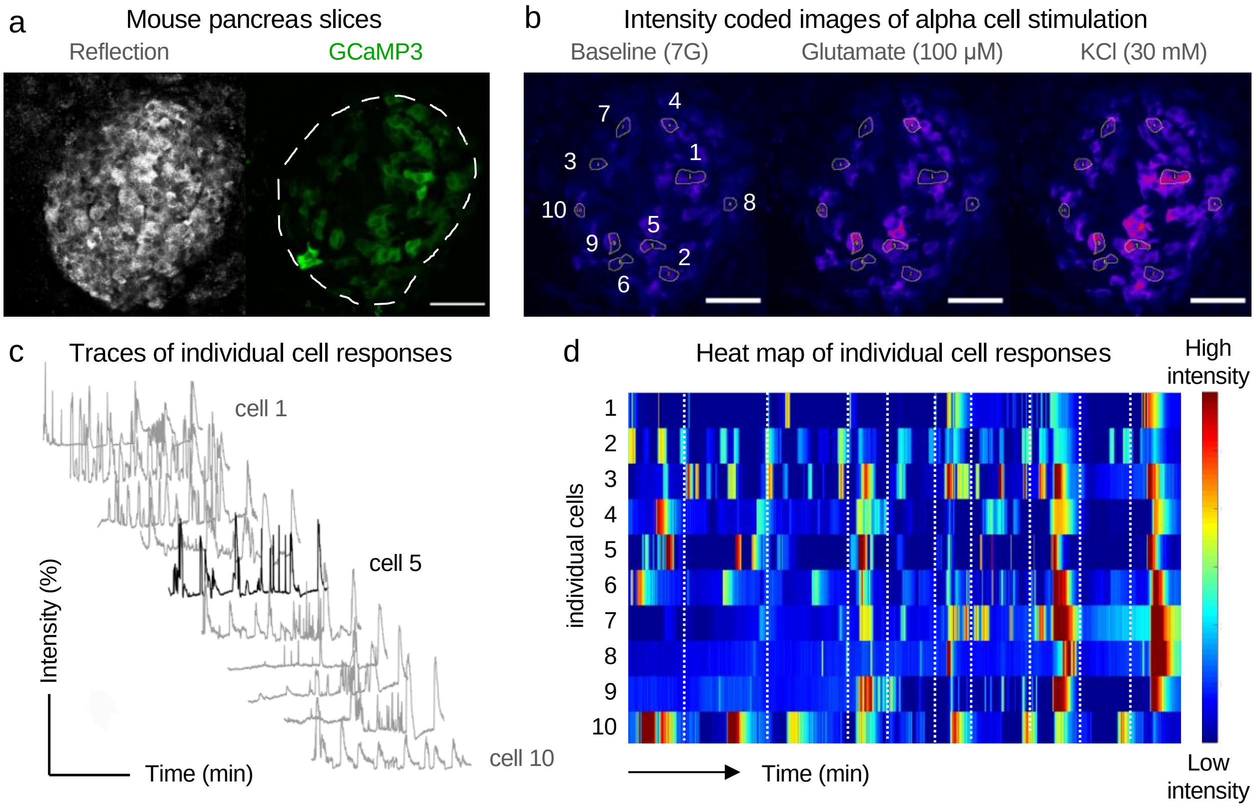


**Figure S2: Detailed description of quantification of cytosolic Ca2$^{+}$ data. Related to Figures 2 and 6.** (a) Z stack of confocal images of an islet within a tissue slice from a Gcg-Cre;GCaMP3 mouse showing reflected light (left) and GCaMP3 signal (right). Reflection is used to identify islets within the slice. (b) Intensity coded z stack images during time of stimulation showing alpha cell activity. Regions of interest were placed around individual alpha cells and changes in fluorescence intensity were expressed as percentage over mean baseline (ΔF/F). Analysis was performed on single planes for each frame. (c) Example traces for marked regions of interest in (b). Due to the vast number of cells average traces were calculated for all cells recorded per animal. Scale bar 50 µm. Dotted line denotes islet border. (d) Heat map showing in vitro $Ca^{2+}$ responses of traces shown in (c). Heat maps are shown to identify specific patterns of individual alpha cells which might get lost in the average traces. Responses are expressed as the fluorescent intensity over baseline of GCaMP3 signal. Each row shows a single cell followed over time in the color scale from low intensity (blue) to high intensity (red) expressed as change in magnitude (%). Dotted line represents stimulus change.

Figure S3

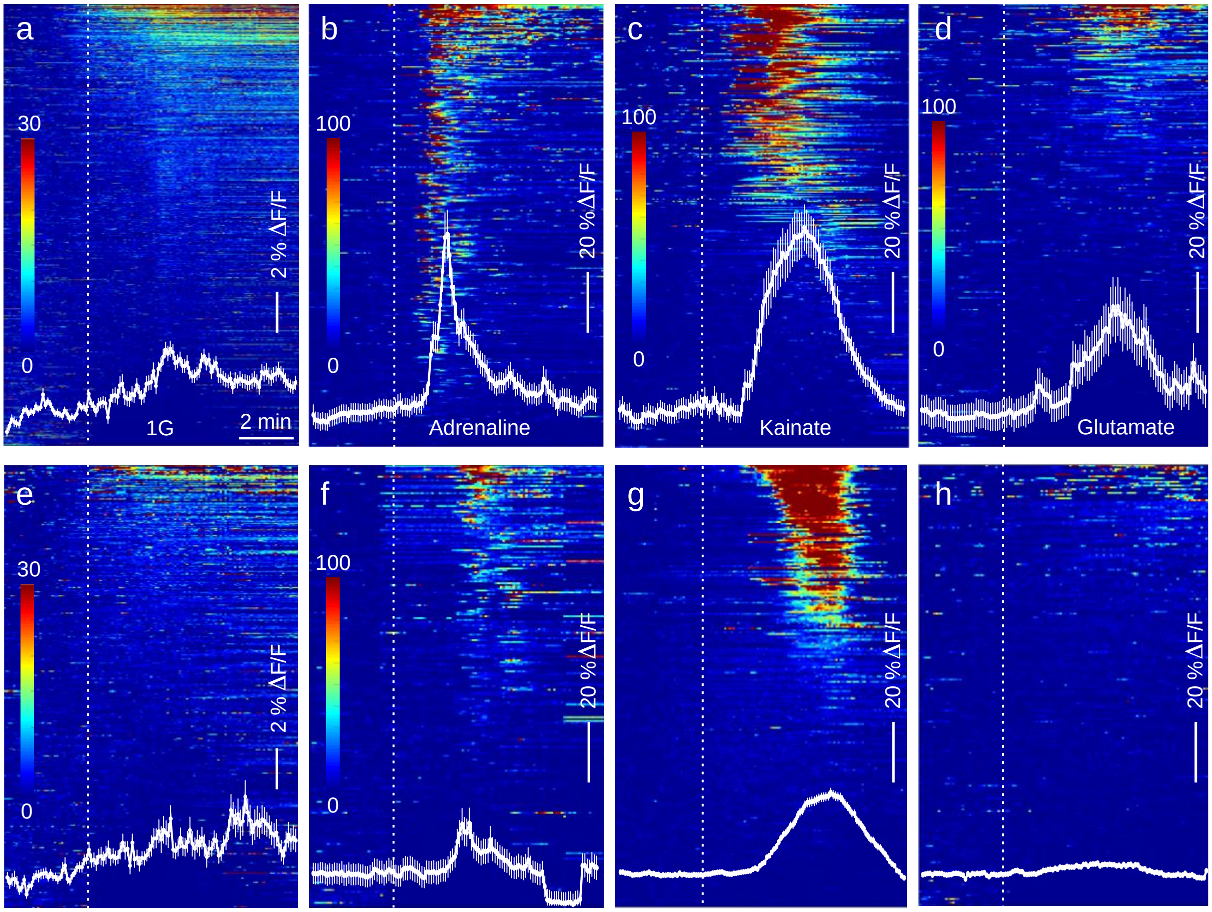


**Figure S3: Defective alpha cell responses to specific stimuli 2 weeks after beta cell ablation. Related to Figure 3.** (a-d) Heat maps showing in vitro $Ca^{2+}$ dynamics of alpha cells within slices from healthy Gcg-Cre;GCaMP3 mice. Responses shown for (a) lowering in glucose concentration from 7 mM to 1 mM, (b) stimulation with adrenaline (10 nM), (c) stimulation with kainate (100 µM) and (d) stimulation with glutamate (100 µM). Average traces of all cells/animal are shown within each heat map. (e-h) Heat maps showing in vitro $Ca^{2+}$ dynamics of alpha cells within slices from rendered diabetic animals 2 weeks after STZ injection. Responses shown for (e) lowering in glucose concentration from 7 mM to 1 mM, (f) stimulation with adrenaline, (g) stimulation with kainate, and (h) stimulation with glutamate. Average traces (± SEM) of all cells are shown within each heat map.

Figure S4

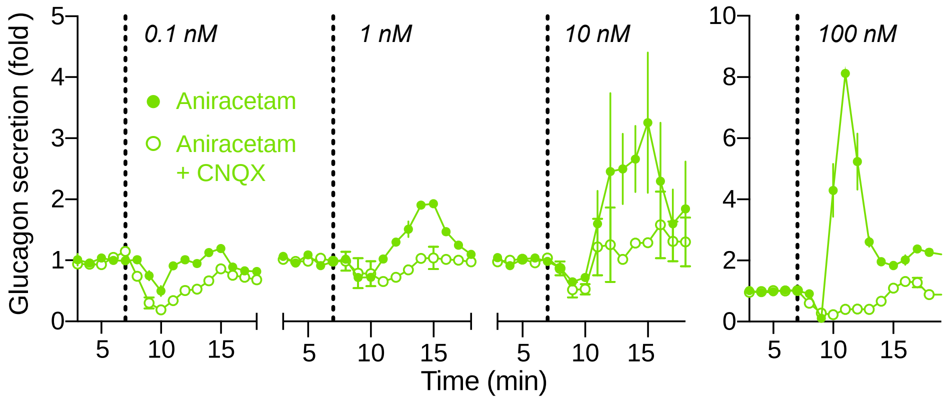


**Figure S4: Concentration response curve of Aniracetam on glucagon secretion in human islets. Related to Figure 5**. Glucagon secretion of isolated human islets (n=3) in response to Aniracetam. Drug concentrations are indicated in every trace. All experiments were performed in HEPES buffer containing 5.5 mM glucose. Traces are normalized to average baseline secretion in the first 5 min and shown as mean (± SEM).

Figure S5

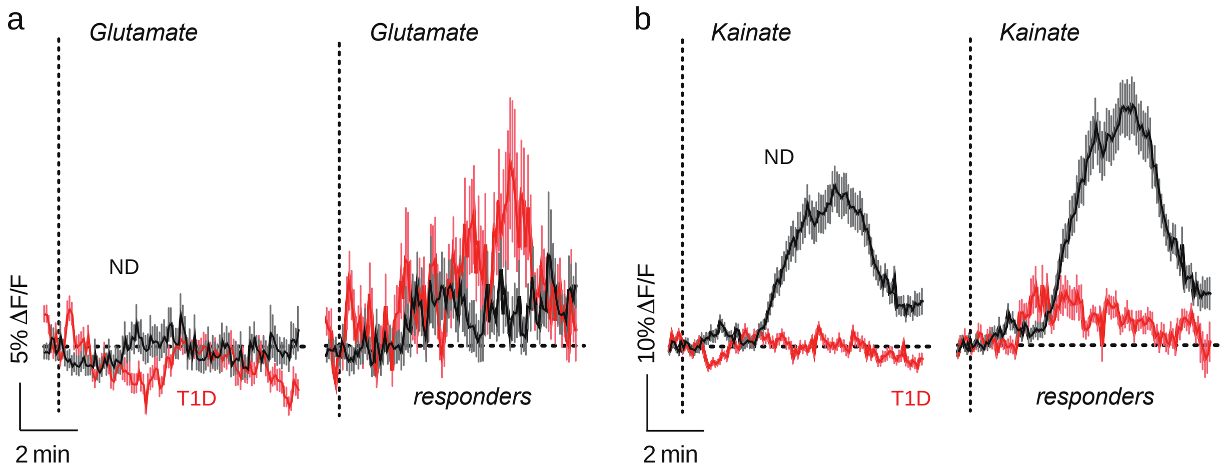


**Figure S5: AMPA/kainate receptor signaling is strongly reduced in type 1 diabetes. Related to Figure 6.** Average traces for $Ca^{2+}$ responses of individual alpha cells from nondiabetic donors (black line) and donors with type 1 diabetes (red line). (a) Average trace of all alpha cells and responders sorted by threshold response to 100 mM glutamate. (b) Average trace of all alpha cells and responders only sorted by threshold response to 100 mM kainate. Traces are shown as mean (± SEM).

Figure S6

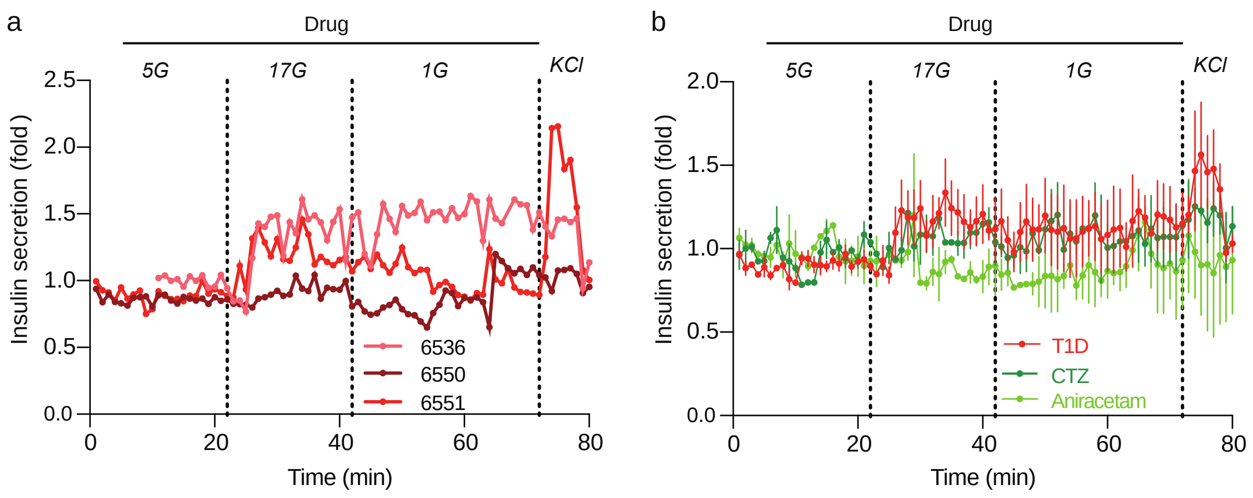


**Figure S6: Positive allosteric modulators aniracetam and cyclothiazide do not have an effect on insulin secretion. Related to Figure 6.** (a) Insulin secretion from tissue slices of 3 individual donors with T1D. Traces are normalized to baseline secretion. (b) Average insulin secretion of the 3 donors shown in (a) in response positive allosteric modulators (100 µM CTZ or 1 mM aniracetam). Traces are normalized to average baseline secretion in the first 5 min and shown as mean (± SEM).
